# Laser induced isolation and cultivation of single microbial cells

**DOI:** 10.1101/2020.07.27.224287

**Authors:** Peng Liang, Huan Wang, Yun Wang, Yinping Zhao, Wei E. Huang, Bei Li

## Abstract

Single cell isolation and cultivation play an important role in studying physiology, gene expression and functions of microorganisms. Laser Induced Forward Transfer Technique (LIFT) has been applied to isolate single cells but the cell viability after sorting is unclear. We demonstrate that a three-layer LIFT system could be applied to isolate single cells of Gram-negative (*E. coli*), Gram-positive (*Lactobacillus rhamnosus* GG, LGG), and eukaryotic microorganisms (*Saccharomyces cerevisiae*) and the sorted single cells were able to be cultured. The experiment results showed that the average cultivation recovery rate of the ejected single cells were 58% for *Saccharomyces cerevisiae*, 22% for *E. coli*, and 74% for *Lactobacillus rhamnosus* GG (LGG). The identities of the cultured cells from single cell sorting were confirmed by using colony PCR with 16S-rRNA for bacteria and large unit rRNA for yeast and subsequent sequencing. This precise sorting and cultivation technique of live single microbial cells can be coupled with other microscopic approaches (e.g. fluorescent and Raman microscopy) to culture single microorganisms with specific functions, revealing their roles in the natural community.

**Importance:** Single cell isolation and cultivation are crucial to recover microorganisms for the study of physiology, gene expression and functions. We developed a laser induced cell sorting technology to precisely isolate single microbial cells from a microscopic slide. More importantly, the isolated single microbial cells are still viable for cultivation. We demonstrate to apply the live sorting method to isolate and cultivate single cells of Gram-negative (*E. coli*), Gram-positive (*Lactobacillus rhamnosus* GG, LGG), and eukaryotic microorganisms (*Saccharomyces cerevisiae*). This precise sorting and cultivation technique can be coupled with other microscopic approaches (e.g. fluorescent and Raman microscopy) to culture specifically targeted single microorganisms from microbial community.

**Abstract Graphic:** 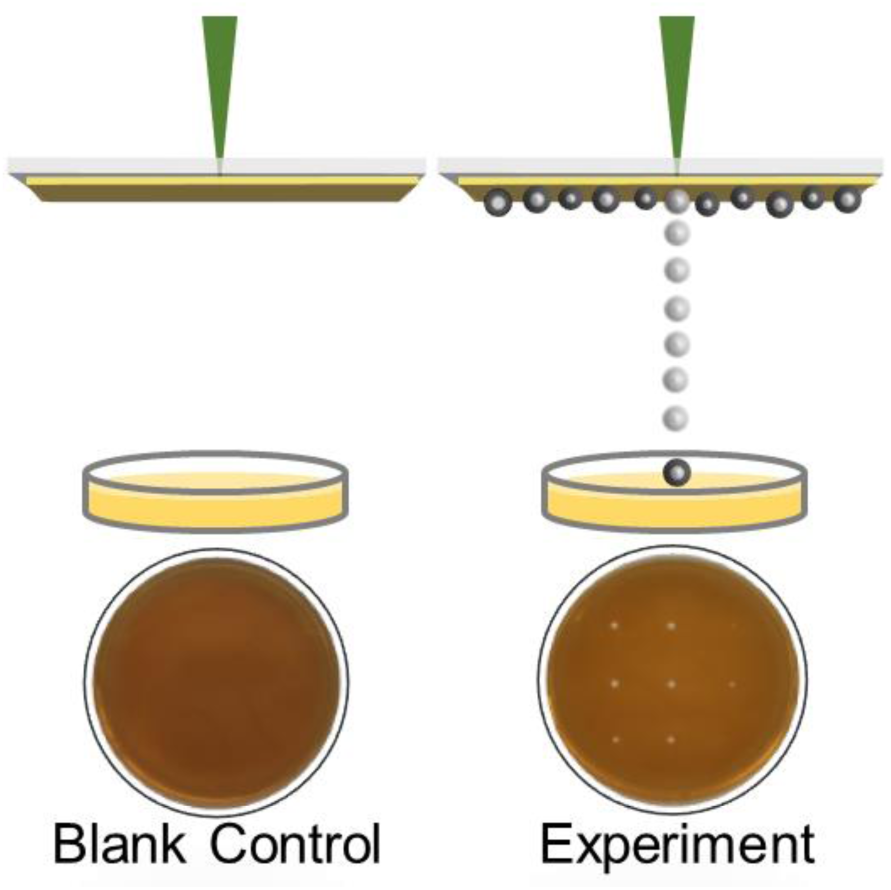

## Introduction

Single cell technology is a powerful tool for studying the growth, physiology, function and biodiversity of microorganisms, especially for the as-yet-unculturable microorganisms in nature (1-5). As the basis and most important step of a single cell study, single cell isolation has been applied in many research fields such as single cell genomics (6), neurobiology (7) and analysis of disease processes (8).

To date, there are a few single cell isolating methods available, such as manual micromanipulation, robotic micromanipulation (9-11), Fluorescence-Activated Cell Sorting (FACS) (12, 13), Magnet-Activated Cell Sorting (MACS) (14, 15) and Laser Microdissection (LMD). Micromanipulation is a commonly used method in the laboratory. However, its throughput is low and requires highly skilled professional training (16). FACS was introduced in 1969 by Leonard Herzenberg (17) and is of wide application with high throughput. FACS requires fluorescent labelling to the targeted cells (13) and lose spatial information of cells due to liquid droplet embedding. Similar with FACS, MACS depends on a magnetic force to isolate cells from the cell suspension in a magnetic field. Although MACS is more rapid and less expensive than FACS, its isolation accuracy is lower than FACS (16). LCM uses a focused laser to cut a cell from its surroundings. It is normally used for fixed tissue, and the viability of the cell is unsure. (18).

Laser induced forward transfer (LIFT) is a promising method for precise single cell isolating (19). LIFT was first described in 1986, when Bohandy et al transferred copper (Cu) onto a silicon substrate using laser irradiation (20). Since then, LIFT has been developed gradually and now is a broadly used printing method which can transfer a great range of materials from electronics to cells, liquid to solid (21). The ejection mechanism of LIFT is dependent on thickness of the coating layer, the material and the laser power and duration, but the basic setup is similar (21). An energy absorption layer is coated on a transparent substrate which we called donor, and then the material to be printed is spread onto the donor. When a laser pulse propagates through the transparent substrate and focuses onto the surface of the energy absorption layer, the energy absorption layer will be heated to vaporize, and the material on it will be pushed away under the gas pressure (22-27) or the shock wave (28, 29) induced by heating.

In the LIFT isolating process, we are able to observe the single cell’ s isolating process under a microscope, and LIFT can also be combined with other optical techniques such as fluorescence imaging (30) and Raman spectroscopy (22-24). LIFT has previously been successfully employed to print mammalian cells including breast cancer cells (31), cardiac cells (32), human osteosarcoma cells (33), neuronal cells, stem cells (34), mouse fibroblast cells (30) and olfactory cells (32), etc. LIFT has been applied for isolation and cultivation of microbial cells as well because of its ability to isolate the bacterial cells without destroying the micro-environment. Haider et al used titanium oxide as the energy absorption layer and studied the effect of laser energy on the viability of yeast and *E. coli* (35). Some other researchers made use of its advantage to isolate and culture the “unculturable” micro-organisms from soil in nature (5) or to analyze the soil microbial community (2). However, Cells usually can not be cultured after LIFT-aided cell ejection (supplementary information Fig. S1). It is a challenge to achieve live cell isolation and subsequent cultivation from single cells.

In this work, we developed a simple three-layer LIFT system to precisely isolate and culture single cells, using typical yeast *Saccharomyces cerevisiae*, Gram-negative bacterium *E. coli*, and Gram-positive bacterium *Lactobacillus rhamnosus* GG (LGG) for the proof of concept.

## Results

### Set up apparatus for single cell sorting

The schematic of the experimental setup is illustrated in Fig. 1a. A 532 nm laser pulse with 5 ns full width half maximum (FWHM) duration (Changchun New Industries Optoelectronics Technology Co., Ltd, China) was utilized for single cell ejection. The laser beam was coupled into a 10× microscope objective (MO1) and focused on the aluminum film (25 nm) coated on the glass. To control the laser pulse, we designed a laser beam expander, a laser energy adjusting module, and a shutter in the optical path (Fig. 1a). The laser beam expander (L1: f=15mm, L2: f=50mm) was used to expand the laser beam from 1 mm to 3.3 mm, which can fit the 10× microscope objective. The laser energy adjusting module contained a half-wave plate (HP) and a polarizing beam splitter (PBS). The laser pulse energy could be changed from 80 nJ to 1300 nJ by adjusting the angle of the half-wave plate (HP). The shutter (S) and laser were programmed to form a single laser pulse. The mirrors (M1-M4) in the setup were used to change the direction of the laser (Fig. 1a).

**Figure 1:**
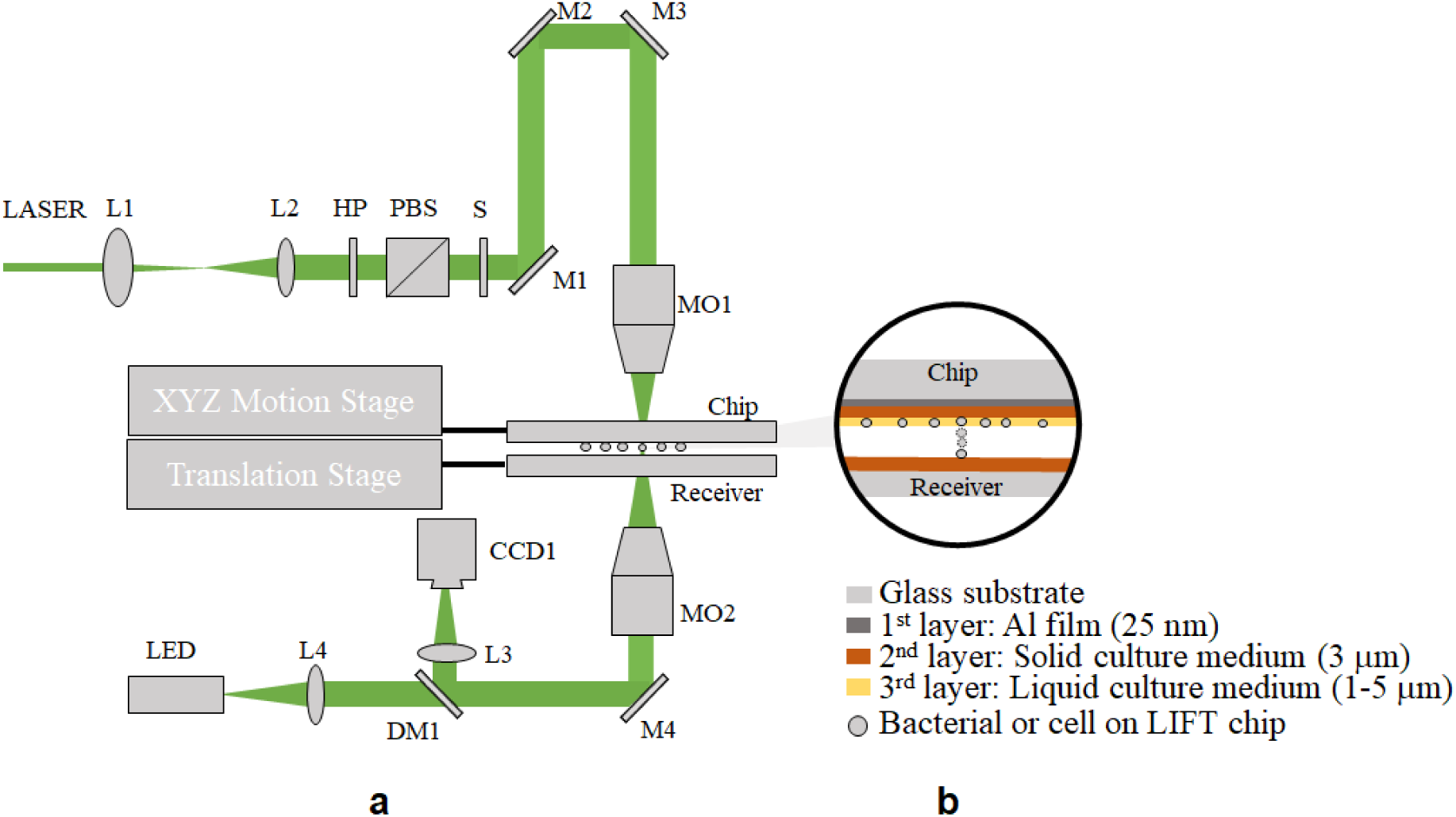
(a) Schematic of the laser induced forward transfer (LIFT) system used for single microorganism isolating and culturing. L1-L4: Lenses; HP: half-waveplate; PBS: Polarizing beam splitter; S: Shutter; M1-M4: Mirrors; DM1: Dichroic mirror; MO1-MO2: Microscopy objectives. (b) Three-layer structure: 1st layer-Al film; 2nd layer-Solid culture medium; 3rd layer-Liquid culture medium.

The cell images were obtained by the bottom imaging system, and the imaging microscope objective (MO2) used here was a 50× Nikon objective. Both the LIFT chip and the receiver were mounted on a translation stage. When cells on the LIFT chip were being examined, the receiver was motored outside of the light path. After a cell was targeted, the receiver would move to the right place to collect the ejected cell.

### Temperature control of the three-layer LIFT system

The chips used for single cell isolation contains three layers as shown in Fig.1b. The first layer (25 nm aluminum film) was used as the energy absorption layer, which absorbs the laser pulse for evaporation, producing a pushing force in an extremely short time. The second layer (agar culture medium) was used as the dynamic release layer (DRL), which lowers down the temperature, protecting the cell from heating damage. The third layer (liquid culture medium) provides a liquid environment for further protection to the target cell from heating (Fig. 1b).

To evaluate this three-layer LIFT system’s ability to protect cells from heating damage, we developed a heat transfer model using a finite element method in the software package Comsol^®^. Fig. 2a shows the schematic of the model. We use three rectangular here to simulate the heat transfer in three layers: the first layer is Aluminum film (25 nm thick), the second layer is Agar (about 3 μm thick) and the third layer is water (3 μm thick, the liquid culture medium).

**Figure 2:**
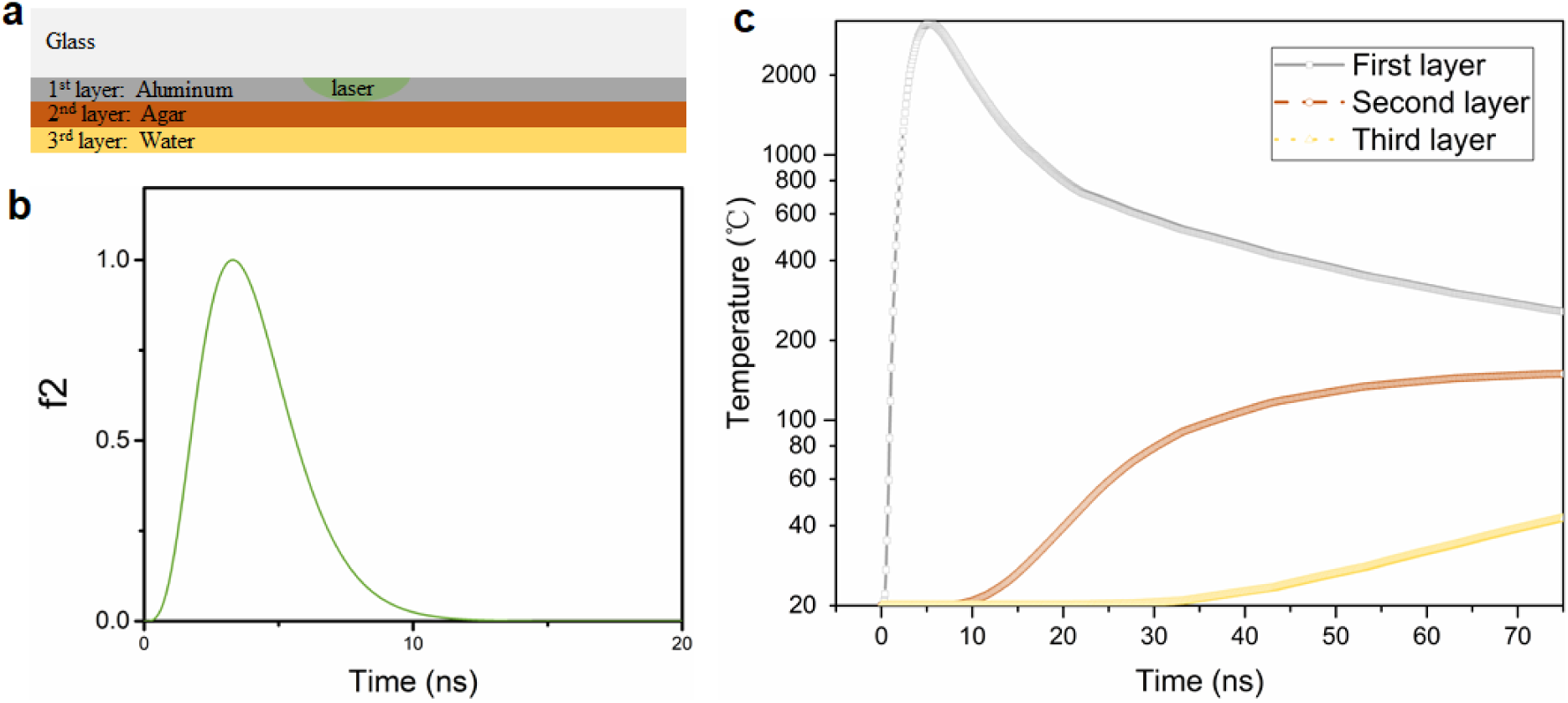
Numerical simulation of the temperature on the surface of each layer with Comsol®. a): Schematic of the FE model. b): Laser pulse’s energy variation along time. The single laser pulse ends at about 10ns. c): Temperature variation of each layer along time.

The governing equation is a time dependent heat transfer equation (Equation 1 and 2). The laser pulse can be considered as a heat source (Equation 3). It has a Gaussian distribution in r direction as written in Equation 4, and the energy variation along time can be written in Equation 5.

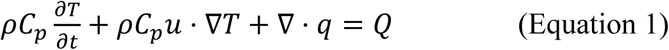

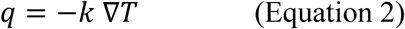

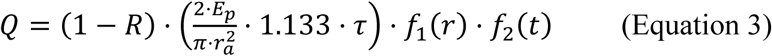

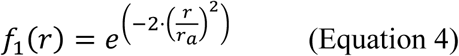

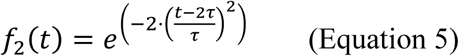

Here, *ρ*: density (*kg/m*^3^); *C*_*p*_: heat capacity (*J/kg. K*); *T*: temperature field (*K*); *t*: time (*s*); *u*: the velocity field (*m/s*); *q*: heat flux (*W/m*^2^); *Q*: heat flux (*W/m*^2^); *k*: thermal conductivity (*W/m. K*); *R*: aluminum film reflectance; *E*_*p*_: laser pulse energy (*J*); *r*_*a*_: beam radius (*m*); *τ*: laser pulse width (*s*).

The physical parameters of each material are listed in Table 1. The density, thermal conductivity and heat capacity are standard properties for aluminum and water. For agar we calculated its density and used the value of thermal conductivity according to a previous report (36). For the heat capacity we used the same value as water. The initial temperature was set as the ambient temperature, 20 °C. A few simplifications and assumptions were introduced to the computation process. First, there are no phase changes in any materials. Second, deformation of the solid culture medium is considered as minimal. Third, the heat can conduct freely between layers if ignoring the thermal contact resistance. The simulation results shown in Fig. 2c suggest that the temperature of the third layer should start rising at about 30 ns. The laser pulse is about 10 ns, and the ejecting process happens at the moment the aluminum’s temperature reaches the evaporating point, which will complete in less than 10 ns. Therefore, theoretically, the cell should have been ejected before the heat is transferred to it. Hence, a cell ejected under this designed system can remain alive.

**Table 1.**
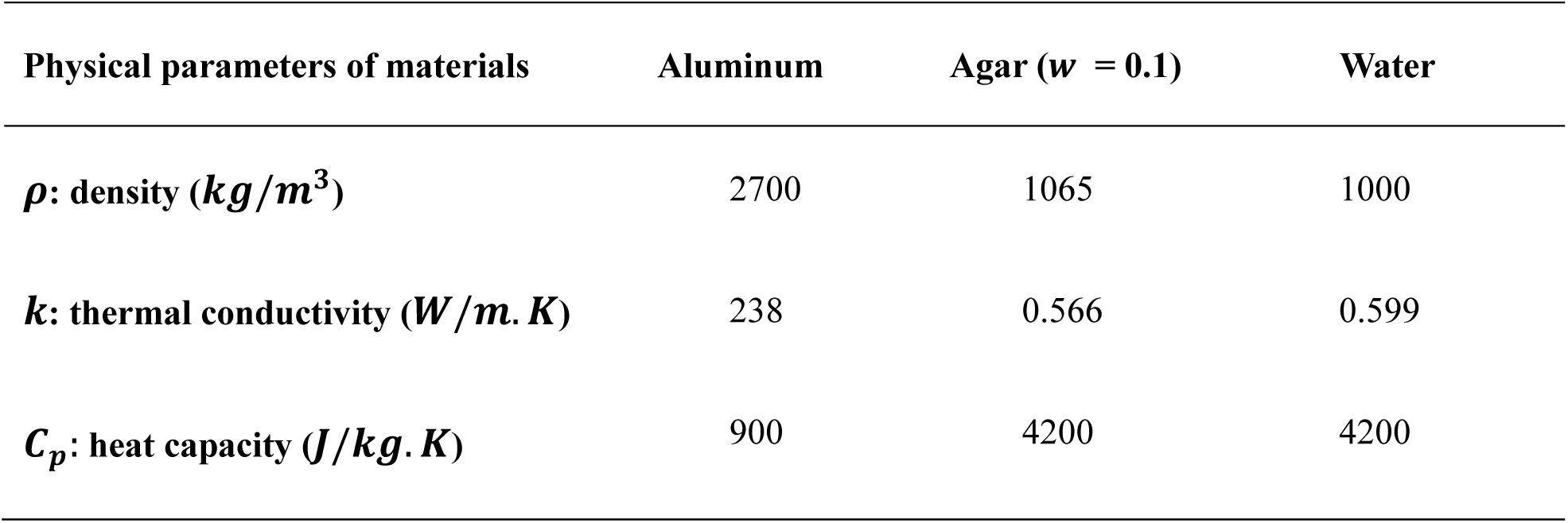
Physical parameters of materials.

| Physical parameters of materials | Aluminum | Agar ($w = 0.1$) | Water |
| --- | --- | --- | --- |
| $\rho$: density ($kg/m^3$) | 2700 | 1065 | 1000 |
| $k$: thermal conductivity ($W/m.K$) | 238 | 0.566 | 0.599 |
| $C_p$: heat capacity ($J/kg.K$) | 900 | 4200 | 4200 |

### Ejecting and capturing of single cells

Precise ejection and collection of single cells is important to single cell isolating and culturing. In this work, *Saccharomyces cerevisiae* and *E. coli* were used to verify the ability of the three-layer LIFT system for isolation and capture of single cells. To image the received cell, we used a 0.17 mm-thick transparent cover slide as the receiver, so individual *Saccharomyces cerevisiae* and *E. coli* cells on the chip and the receiver could be clearly observed by adjusting the bottom objective to focus on different planes, and the distance between the chip and receiver is about 250 μm. Figure 3 demonstrates that single *Saccharomyces cerevisiae* and *E. coli* cells were ejected from the chip, leaving a mark on the chip, and the ejected cells were able to be retrieved on the receiver. When five single cells of *Saccharomyces cerevisiae* and *E. coli* were ejected, we could find five single cells on the receiver afterwards (Fig.3). The distribution of the ejected cells on the receiver is even similar to that originally on the chip (Fig.3, panel m-p). It needs to be mentioned that we also received five single *Saccharomyces cerevisiae* cells on the receiver after ejecting five individual *Saccharomyces cerevisiae* cells (Fig.3, panel e-h), but one of them was out of the viewing field (data not shown). These results suggested that the single cell sorting system could precisely isolate individual yeast and bacterial cells.

**Figure 3:**
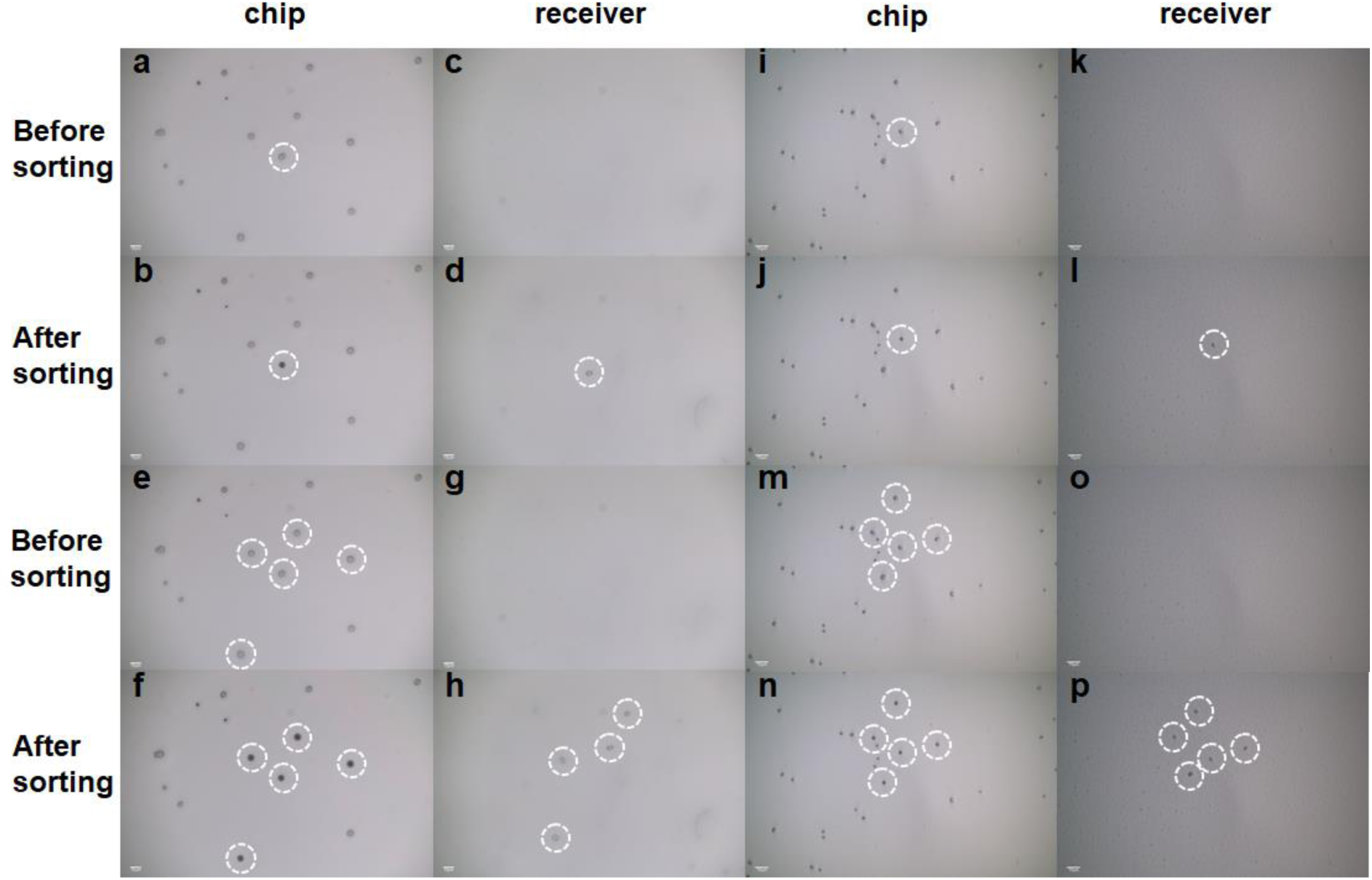
Isolating and receiving single *S. cerevisiae* and *E. coli* cells one by one, the bar represents 20 μm. Panels a-d show the results of one single *S. cerevisiae* cell isolating and receiving. a) Chip before sorting. b) Chip after sorting. c) Receiving slide before sorting. d) Receiving slide after sorting. Panel e-h show the results of isolating 5 *S. cerevisiae* cells and receiving. e) Chip before sorting f) Chip after sorting. g) Receiving slide before sorting. h) Receiving slide after sorting. Note: Only 4 *S. cerevisiae* cells were viewed in panel h. Actually, the fifth cell was also collected, but it was not in the same field of view as these four cells under the microscope. Panel i-l show the results of one single *E. coli* isolating and receiving. i) Chip before sorting. j) Chip after sorting. k) Receiving slide before sorting. l) Receiving slide after sorting. Panel m-p show the results of isolating 5 *E. coli* cells and receiving 5 *E. coli* cells. m) Chip before sorting. n) Chip after sorting. o) Receiving slide before sorting. p) Receiving slide after sorting.

### Sorting single cells for cultivation using the ejection system

*Saccharomyces cerevisiae, E. coli* and *Lactobacillus rhamnosus* GG (LGG) were chosen as the representative eukaryotic and prokaryotic Gram-negative and Gram-positive cells for the single cell sorting and cultivation in this study. The sorting system was placed inside a clean lamina flow hood. The control group with the blank area near the targeted cells ejected did not lead to any colony growth on the receiving agar plates (Fig. 4, the first row). A single cell sorting of *Saccharomyces cerevisiae, E. coli* and LGG formed one single colony on the corresponding agar plates (Fig. 4 the second row), demonstrating that the sorted single cells by ejection was still alive and able to form single colonies. This shows that singe cell isolating and culturing could be indeed achieved using this three-layer LIFT system.

**Figure 4.**
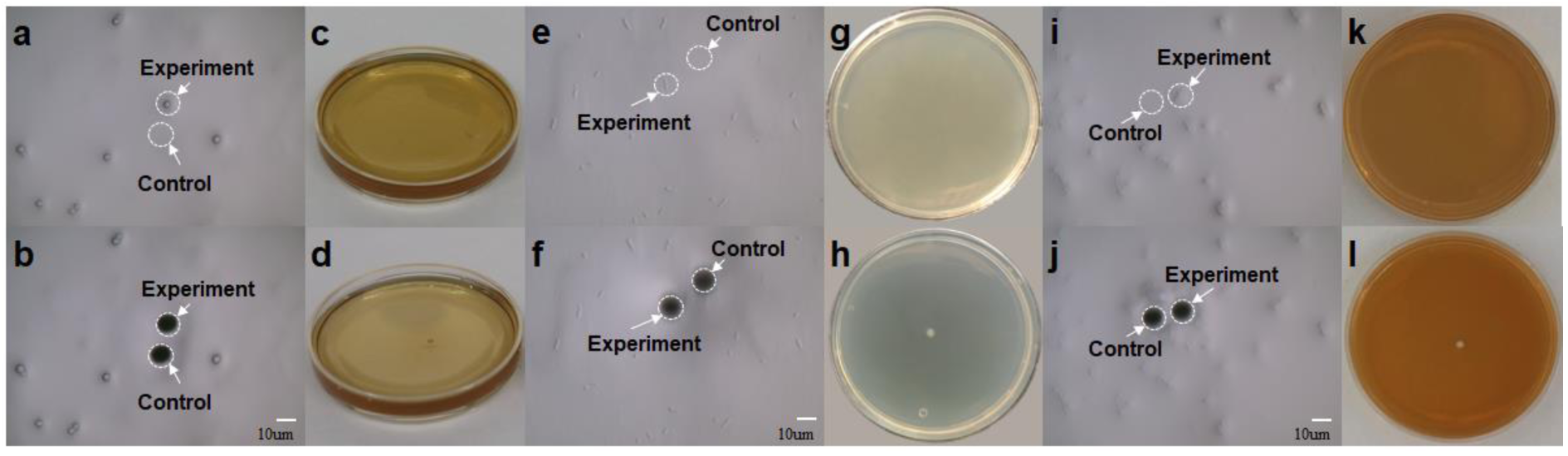
Ejecting and culturing results of single cells of *S. cerevisiae, E. coli* and *Lactobacillus rhamnosus* GG (LGG). Panels a-d show the results of sorting one single *S. cerevisiae* cell and culturing for 36 h. a) Chip before sorting. b) Chip after sorting. c) Results of the control group after culturing 36 h. d) Results of the experiment group after culturing 36 h. Panels e-h show the results of sorting one single *E. coli* cell and culturing for 16 h. e) Chip before sorting. f) Chip after sorting. g) Results of the control group after culturing 16h. h) Results of the experiment group after culturing 16 h. Panels i-l show the results of sorting one single LGG cell and culturing for 48 h. i) Chip before sorting. j) Chip after sorting. k) Results of the control group after culturing 48 h. l) Results of the experiment group after culturing 48 h.

Fig 5 shows that ejection of *Saccharomyces cerevisiae* at 9 different receiving places in one petri dish (Fig 5a) by controlling the translation stage, and 1, 2 or 10 individual *S. cerevisiae* cells were ejected respectively in each area (Fig 5b). Five replicates were carried out in each case. The average recovery rate of single yeast cell ejection could reach 58%, and the recovery rates were 69% and 71% for 2 and 10 yeast cell sorting respectively (Fig. 5 and Table 2).

**Table 2.**
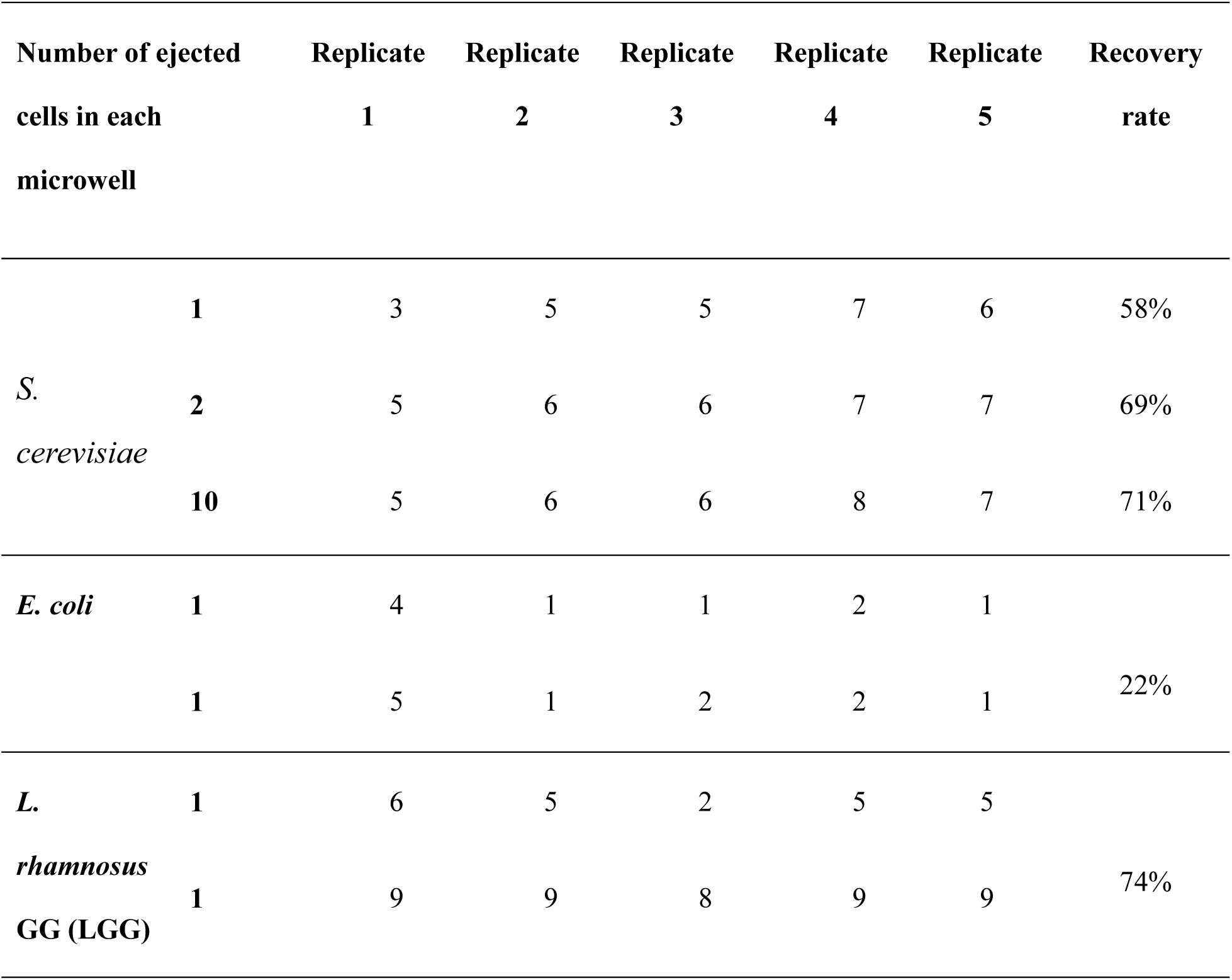
Statistics of *S. cerevisiae, E. coli* and LGG’s culturing results.

| Number of ejected<br>cells in each<br>microwell |  | Replicate<br>1 | Replicate<br>2 | Replicate<br>3 | Replicate<br>4 | Replicate<br>5 | Recovery<br>rate |
| --- | --- | --- | --- | --- | --- | --- | --- |
| *S.*<br>*cerevisiae* | 1 | 3 | 5 | 5 | 7 | 6 | 58% |
|  | 2 | 5 | 6 | 6 | 7 | 7 | 69% |
|  | 10 | 5 | 6 | 6 | 8 | 7 | 71% |
| *E. coli* | 1 | 4 | 1 | 1 | 2 | 1 | 22% |
|  | 1 | 5 | 1 | 2 | 2 | 1 |  |
| *L.*<br>*rhamnosus*<br>GG (LGG) | 1 | 6 | 5 | 2 | 5 | 5 | 74% |
|  | 1 | 9 | 9 | 8 | 9 | 9 |  |

**Figure 5.**
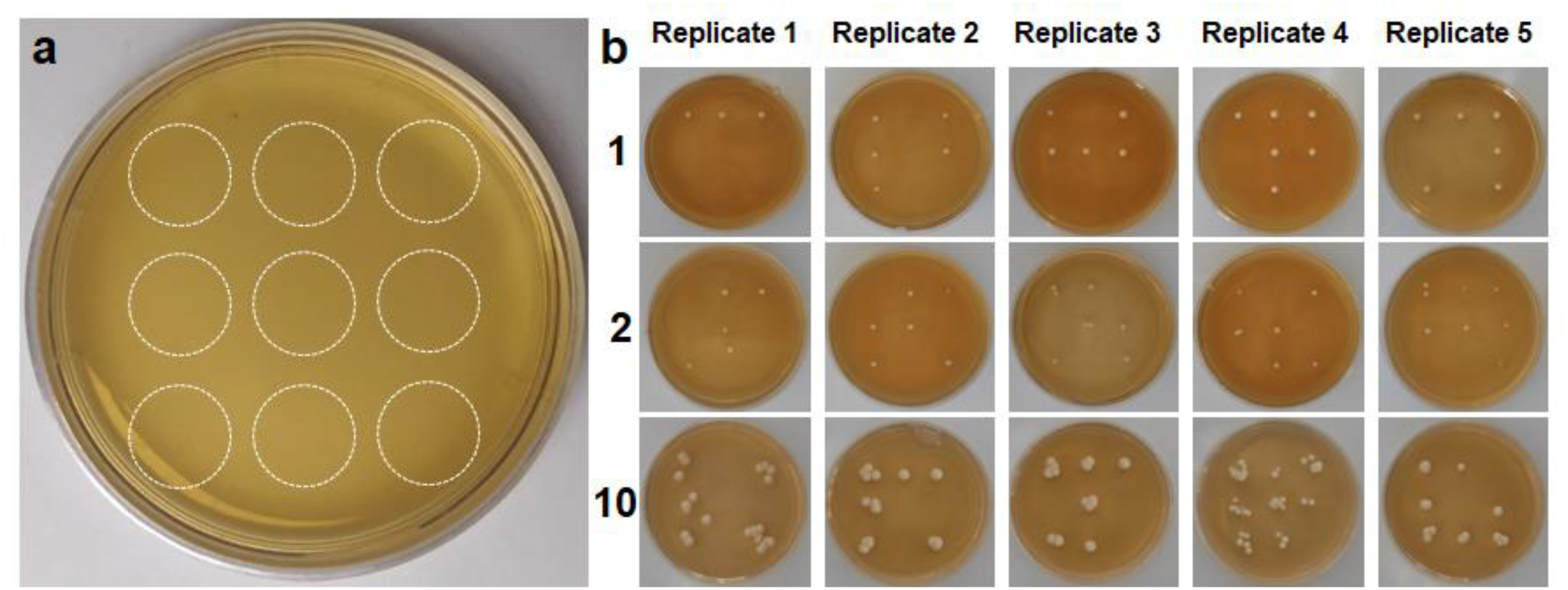
The results of culturing ratio of *S. cerevisiae* cell isolating and culturing. a) 9 receiving positions in one petri dish. b) The results of ejecting 1, 2, 10 single *S. cerevisiae* cells in each hole.

We have also ejected one single *E. coli* and LGG bacterial cells and retrieved their viability and cultivability by counting the number of colonies (Fig 6). Ten replicates were performed in each case. It shows that the recovery rates of single *E. coli* and LGG ejection were 14% and 74% respectively (Fig. 6 and Table 2). It is likely that Gram-positive LGG was robust to survive after ejection sorting due to its thick cell wall.

**Figure 6.**
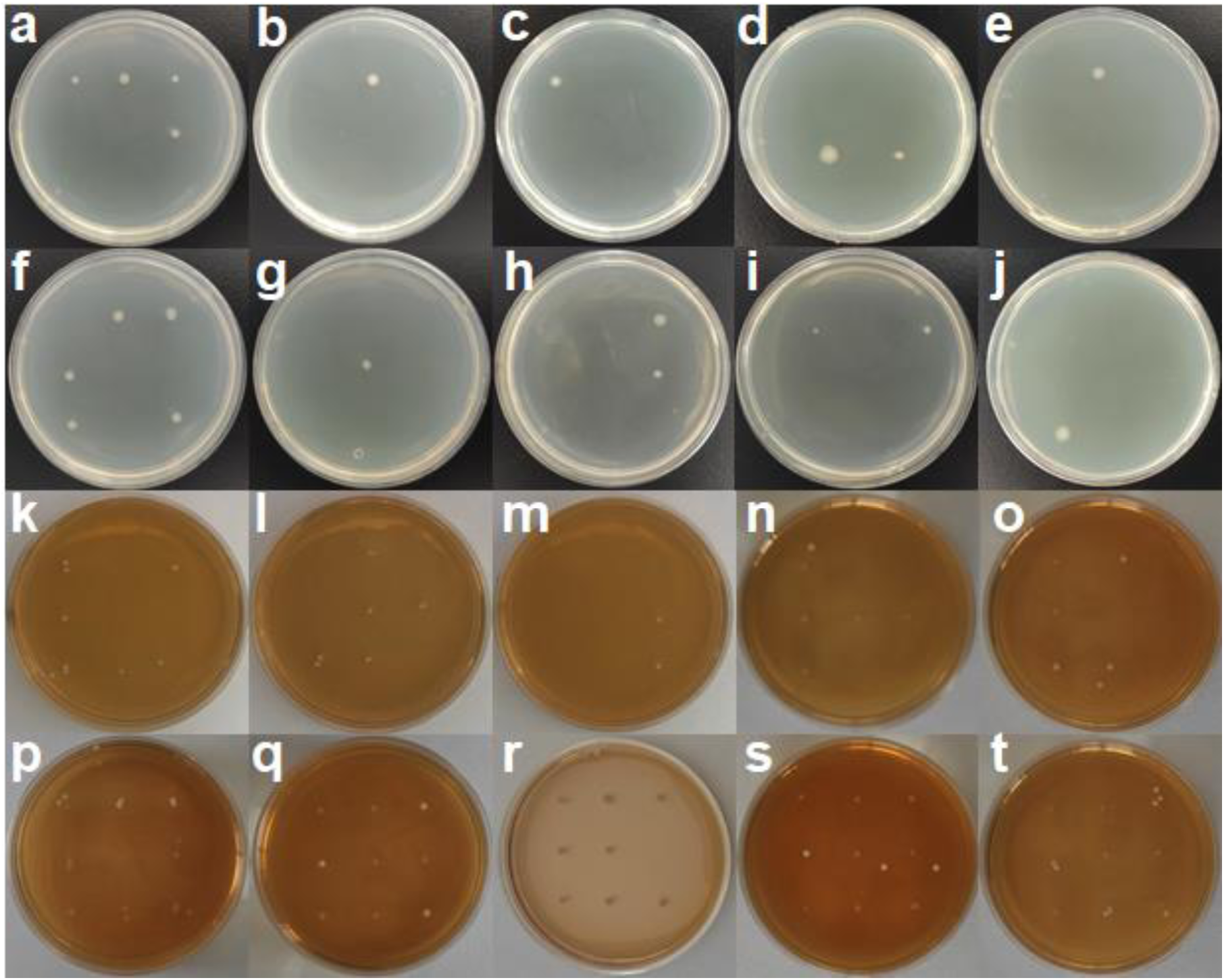
Culturing results of isolated single *E. coli* and *Lactobacillus rhamnosus* GG (LGG) cells in each receiving position. Panel a-j ejecting one single *E. coli* cell in each hole and culturing for 16 h. Panels k-t ejecting one single LGG cell in each hole and culturing for 48 h.

To verify the sorted single cells were the originally targeted cells, five colonies of each species were randomly selected for colony PCR using yeast 18S (for *Saccharomyces cerevisiae*) and universal 16S rRNA primers (*E. coli* and LGG) respectively. The PCR products were purified and sequenced (Supplementary information Fig. S2). The sequencing results confirmed that the sorted cells were the original target cells (Supplementary information Fig. S3)

The results demonstrate that single yeast and bacterial cells can be sorted by the LIFT ejection system while remaining alive to be able to form colonies.

## Discussion

### Precise and accurate single-cell ejecting and capturing

In this study, single cell ejecting, capturing (Fig.3) and culturing (Fig.4, Fig.5 and Fig.6) have been accomplished by using a three-layer LIFT system. Although LIFT has been reported in printing a great range of materials from electronics to cells, liquid to solid, it has not been easily applied in accurate single cell ejecting and culturing because of the heat and force in the LIFT process. Many researches focused on printing mammal cells from liquid layer, but a microbial cell is smaller than a mammal cell. We need the liquid layer to be thin enough (thinner than the diameter of the *Saccharomyces cerevisiae* or *E. coli*), so that the *Saccharomyces cerevisiae* or *E. coli* cells cannot move freely in the liquid layer and we are able to focus on the target cell.

When doing experiments, the environment conditions such as the platform vibration, air flow or other micro-environments could affect the success rate of cell ejecting, receiving and culturing. These factors can mostly be avoided by placing the experiment setup on an active vibration isolation optical table sealed with a closed cover. Besides, the laser spot’s position and energy also need to be carefully adjusted. If the spot deviates too far from the target or the energy is not moderate, three cases may happen. Firstly, the cell may not be successfully ejected. Secondly, the ejected cell flies away and does not land on the right receiving position. Finally, other cells around the target cell are also ejected and the number of the received cell is more than planned or contamination occurs. To address above issues, we use a 500 nJ laser pulse to eject the *Saccharomyces cerevisiae* and 300 nJ for *E. coli* and *Lactobacillus rhamnosus* GG (LGG). The effect of the laser pulse energy, the agar’s thickness and other materials can be studied further in the future.

Compared to other single cell isolation methods we mentioned before, LIFT overcomes the shortcomings existed in those methods: (1) Need to label the cells with fluorescent probes or magnetic beads, and the cell type is limited. (2) Need highly-time-consuming operation, or costs. (3) Need a high quantity of cell suspension. (4) The viability of the isolated single cell is not good enough for cultivation.

### Single cell viability and cultivability

Laser radiation, heat and force in the ejecting, flying and landing processes, and dry environment could cause photo- or thermo-damage to single cells in the LIFT process. The damage could be reduced by the three-layer design. The impact of laser radiation on cellular viability is negligible because only a tiny amount (0.1%) of the laser pulse can pass through the Aluminum film. As the intermediate layer, solid culture medium cannot only protect the cells from damage of high temperature and pushing force of LIFT, but also slows down the drying speed of liquid and provides nutrition to the cell in a short time even if the liquid dries out.

Although other adiabatic materials such as polyimide have been used as the intermediate layer, agar medium is still a good choice because it is very cheap and easy to obtain. Furthermore, agar is not as strong as polyimide, so it needs less energy to achieve single cell transfer, especially a microbial cell. We used only a 500 nJ laser pulse for *Saccharomyces cerevisiae* and 300 nJ for *E. coli* and LGG in the experiment.

Although agar can slow down the liquid’s drying speed, it would still dry out after long time plating onto the chip. We think a dry environment kills the cells in a short period. In another experiment (not included in this report), we get the conclusion that 50% yeast cells died 30 minutes after drying on the three-layer LIFT chip by staining with methyl blue. So next step, we will develop new methods to further slowdown the liquid’s drying speed to keep the cell’s viability.

To our knowledge, this is the first report that accomplishes one single bacterial or yeast cell isolation and culturing using LIFT, and the average recultivation ratio of the received single cells reaches 58%, 22%, 74% for *Saccharomyces cerevisiae, E. coli*, and *Lactobacillus rhamnosus* GG (LGG) respectively by using the three-layer LIFT system. The results demonstrate that the three-layer LIFT system can not only reduce heat damage to the cells in the ejecting process but also provide a long-duration liquid environment. Therefore, it is a promising method to isolate and culture the “unculturable” microorganisms existing in the natural habitat such as soil or ocean sediments.

## Materials and Methods

### LIFT chip and receiver

For the LIFT chip, we proposed a three-layer design. A 25-nm thickness aluminum film was used as the first layer and was coated on a glass slide. The aluminum film absorbs the laser pulse and will be heated and form the ejecting force. Then an agar medium (YPD agar for *Saccharomyces cerevisiae*, MRS agar for LGG, LB agar for *E. coli*) (5g Broth and 1.5g agar in 100mL water) film was spread as the second layer by a spreading machine. To make the solid culture medium film more uniform, three PA (Phytic Acid)-Al^3+^ films was attached onto the surface of the aluminum film to make it more hydrophilic(37). The procedure was performed as follows: the cleaned chips were soaked in 0.255 mmol/L PA solution for 10 minutes and then soaked in 55.5 mmol/L AlCl_3_ solution for 2minutes, rinsed with ultra-pure water and dried with a rubber suction bulb. This process was repeated 3 times to obtain 3 layers of PA-Al^3+^ complex on the chip. Then we heated the solid culture medium to liquid state in an oven, and spread it onto the surface of the chip with a spreading machine (KW-4A, Institute of Microelectronics of the Chinese Academy of Sciences) rotating at a speed of 500 rpm for 16 seconds and 100 rpm for 2 seconds. The thickness of the solid culture medium is roughly estimated as 5 μm under a microscope. After the LIFT chip was cooled in a 4°C refrigerator for a while, we directly spread the yeast cell solution or bacteria solution (the third layer) on the LIFT chip, and then clamped it by a device which was mounted on a XYZ motion stage for further observation and selecting under a microscope.

For the receiver, a 35 mm petri dish (Thermal Fisher) with agar medium was used and fixed on a device which is mounted on a translation stage. The distance between the receiver and the donor is about 2 mm. Besides, we wrote a program to control the translation stage to stop at 9 different points corresponding to the 9 receiving points on the petri dish.

### Cell cultivation, laser-induced transfer of single cell, and recovery of sorted cells

A single colony of Saccharomyces cerevisiae, *E.coli* or LGG was inoculated into 10 ml YPD, LB or MRS Broth respectively, and cultured in a 30°C or 37°C incubator for 16 hours at a shaking speed of 200 rpm till an OD600 of 1.8, to mention, LGG was cultured in 37°C incubator not shaking.

For the LIFT process of single cell ejecting and culturing, different laser energy were applied for the three species, 300 nJ for *E. coli* and LGG, 500 nJ for *Saccharomyces cerevisiae* Yeast. By controlling the upper XY motorized stage (Fig.1), we selected the target cell and put it under the laser pulse, and then the receiver reached under the target cell. After ejection, the receiver was moved out of the optical path. To avoid contamination, the experiment set-up and the chips and receiver were all put into a laminar flow hood which was exposed to UV light for 20 minutes before doing experiment.

After the LIFT single cell ejecting process, the receiving petri dishes were put in a 30 °C incubator and cultured for about 36 hours for *Saccharomyces cerevisiae*, or a 37 °C incubator for 16 hours for *E. coli*, and a 37 °C incubator for 48 hours for LGG.

### Colony PCR and sequencing

For *Saccharomyces cerevisiae*, 5 colonies from the cultivation plate were randomly picked and mixed into 500 μL ultra-pure water. Then 2 μL of this solution, 10 μL Taq, 1 μL NL1 primer (10 μM concentration), 1μL NL4 primer (10 μM concentration) (38, 39), and 6 μL DEPC-treated water (R1600, Solarbio) were mixed together (20 μL) into a PCR tube. For *E. coli* and *Lactobacillus rhamnosus* GG (LGG), 5 colonies from the cultivation plate were randomly picked and mixed into 1 mL ultra-pure water respectively, and then 2 μL of the solution, 10 μL Taq, 1 μL 27F primer (10uM concentration), 1μL 1492R primer (10 μM concentration) (38), and 6 μL DEPC-treated water (R1600,Solarbio) were mixed together as the PCRs (20 μL).

The PCR reaction was performed on a T100(tm) Thermal Cycler (Bio-Rad). The amplification program used here was as follows: 95 °C for 3 min; 35 cycles of 95 °C for 15 s, 55 °C for 30 s, and 72 °C for 1 min; and 72°C for 5 min. The Gel electrophoresis experiment was performed on PowerPac(tm) (Bio-Rad) at 130V for 25 min and the sequencing process was performed by Sangon Biotech.

## Acknowledgements

We thank Hooke Instruments Ltd. for finance support (PHIB2004-005) and the help on biological experiment details.

## Competing interest

The authors declare that they have no competing interests.

